# Sequential cold and heat stresses establish an intergenerational stress memory in rapeseed (*Brassica napus* L.)

**DOI:** 10.1101/2025.07.24.666528

**Authors:** İrem Çağlı, Aylin Gazdağlı Talay, Mine Berrak Halik, Fatma Tunalı, Çağla Sönmez

**Affiliations:** Department of Molecular Biology and Genetics, Necmettin Erbakan University, 42090 Konya, Türkiye; Department of Molecular Biology and Genetics, Faculty of Agriculture and Natural Sciences, Konya Food and Agriculture University, 42080 Konya, Türkiye; Department of Biotechnology, Konya Food and Agriculture University, Konya 42080, Türkiye; Botanic Garden “Giardino dei Semplici”, Department of Pharmacy, “G. d’Annunzio” University, 66100, Chieti, Italy; Department of Medical Biology, School of Medicine, Atilim University, 06830 Ankara, Türkiye

**Keywords:** *Brassica napus*, Temperature stress, Cross-tolerance, Priming, Intergenerational stress memory

## Abstract

Plants frequently experience temperature extremes that threaten growth and reproduction, yet their ability to retain and transmit stress responses across generations remains poorly understood. In this study, we investigated whether early cold exposure primes rapeseed seedlings for enhanced heat tolerance and whether such effects are inherited by the next generation. Seedlings were subjected to cold stress (4 °C for 3 weeks), heat stress (38 °C for 2 days), or sequential cold followed by heat stress. Control plants were grown under optimal conditions. We evaluated physiological, biochemical, and molecular traits in both the treated plants and their first-generation progeny.

Temperature stress influenced flowering time, seed weight, seed oil content, and fatty acid composition. Genes involved in fatty acid metabolism, including *BnaFAD2*, *BnaFAD5*, *BnaFATB*, and *BnaWD40*, were differentially expressed. In the progeny of sequentially stressed plants, total phenolics, flavonoids, antioxidant activity, and chlorophyll content were significantly elevated, indicating the presence of intergenerational stress memory.

Our findings show that sequential cold–heat stress not only enhances immediate stress tolerance but also induces heritable metabolic and physiological adaptations. These results provide new insights into the mechanisms of cross-tolerance and the potential for exploiting intergenerational stress memory in crop improvement.

## Introduction

Plants are frequently exposed to environmental stress throughout their life cycle. The intensity of abiotic stresses such as extreme temperatures, drought, and salinity is increasing due to climate change, posing major threats to crop productivity and global food security (Benitez-Alfonso et al., 2023). As a result, enhancing stress resilience has become critical for sustainable agriculture.

Stress resilience relies on a plant’s ability to adapt at molecular and physiological levels to maintain cellular integrity and survive adverse conditions (Benitez-Alfonso et al., 2023). In temperate regions, crops like rapeseed (*Brassica napus* L.) are particularly vulnerable to temperature extremes, including cold and heat stresses (Zajac et al., 2016). Cold stress can occur as either freezing (< 0°C) or chilling (0–20 °C), lasting from hours to weeks (Guo et al., 2018). Prolonged cold, or vernalization, promotes timely flowering in many plants (Henderson et al., 2003).

Cold stress disrupts key physiological processes, such as enzyme activity, metabolite transport, and membrane fluidity (Thomashow, 1999; van Buer et al., 2016). These effects are initiated by early sensing mechanisms, including calcium signaling and transcription factor activation, which trigger adaptive gene expression (Thomashow, 1999; Byun et al., 2014; van Buer et al., 2016). Cold tolerance involves activation of antioxidative systems, particularly in chloroplasts and mitochondria, the primary sources of reactive oxygen species (ROS) (Byun et al., 2014; Li et al., 2014). ROS initially act as signals to activate stress-responsive genes, including those encoding heat-shock proteins and enzymes related to osmolyte and hormone metabolism (Hossain et al., 2015; Hossain et al., 2018). As stress persists, ROS are counteracted by upregulated antioxidant pathways (Almeselmani et al., 2006; Hossain et al., 2015; Liu et al., 2022).

Maintaining membrane integrity is essential for cold tolerance, and changes in lipid composition—particularly fatty acid unsaturation—are closely linked to stress adaptation (Miquel et al., 1993; Gao et al., 2024). In a recent study, we showed that prolonged cold exposure during early development leads to altered seed lipid composition in mature rapeseed plants (Çağlı et al., 2025). This suggests that cold-induced changes in lipid metabolism may be retained through reproductive development, even in the absence of ongoing stress. The epigenetic memory of vernalization and its link to flowering time is well documented in Arabidopsis (De Lucia et al., 2008; Menon et al., 2021). However, whether cold-induced changes in seed lipid metabolism represent another form of epigenetic memory remains unclear.

The effects of heat stress—another major temperature-related challenge—have been extensively studied and reviewed (Kan et al., 2023; Distéfano et al., 2025). Like cold, heat stress disrupts membrane integrity, impairs the photosynthetic apparatus, and increases ROS production, leading to oxidative damage (Huang et al., 2019; Rashid et al., 2020; Kan et al., 2023; Distéfano et al., 2025). Consequently, plants initiate similar molecular and physiological responses under both cold and heat stress (Hossain et al., 2018; Kan et al., 2023; Distéfano et al., 2025). Rapeseed is particularly sensitive to heat stress during reproductive stages, where seed yield, oil content, and composition are significantly affected (Baux et al., 2008; Baux et al., 2013; Yu et al., 2014; Rahaman et al., 2018; Huang et al., 2019; de Almeida et al., 2021; Secchi et al., 2023). However, the effects of early heat exposure before floral induction remain largely unexplored.

Studies have shown that an initial, mild stress can enhance a plant’s ability to withstand subsequent, more severe stress—a phenomenon known as cross-tolerance (Fu et al., 1998; Freidrich et al., 2019; Liu et al., 2022). Cold priming has been reported to improve heat tolerance in various species, including winter rye, grape, tomato, and barley (Fu et al., 1998; Wan et al., 2009; Mei & Song, 2010; Li et al., 2014). Cross-tolerance is often mediated by priming, which allows plants to establish stress memory. This memory may be short-term (lasting days) or long-term (across generations), involving transcriptional regulation and epigenetic changes (Lämke & Bäurle, 2017; Friedrich et al., 2019; Oberkofler et al., 2021; Liu et al., 2022). ROS signaling and antioxidant defenses play central roles in this process (Mei & Song, 2010; Li et al., 2014; Hossain et al., 2015; Hossain et al., 2018; Oberkofler et al., 2021; Liu et al., 2022).

While much is known about immediate stress responses, less attention has been paid to the intergenerational effects of stress. The ability of plants to pass on stress memory to progeny— thereby enhancing resilience—is an emerging area of research. Intergenerational stress memory refers to inherited changes observed in the immediate next (stress-free) generation, whereas transgenerational memory extends across at least two successive stress-free generations (Oberkofler et al., 2021). Temperature-related transgenerational memory has been shown in Arabidopsis and wheat (Whittle et al., 2009; Suter & Widmer, 2013; Wang et al., 2016; Groot et al., 2017). In rapeseed, drought stress has been reported to enhance seed vigor in the next generation (Hatzig et al., 2018). However, to our knowledge, the inter- or transgenerational effects of sequential cold and heat stresses have not yet been investigated.

This study aims to determine whether cold stress during the seedling stage can prime rapeseed plants for enhanced heat tolerance later in development and whether such sequential exposure induces heritable changes. Specifically, we investigated whether early cold priming triggers physiological, biochemical, and molecular responses that improve heat stress tolerance during seed development, and whether these responses persist in the next generation. By analyzing seed and seedling traits, antioxidant capacity, and fatty acid metabolism gene expression, we explore the potential for cold-induced stress memory and its intergenerational transmission in rapeseed. These findings offer new insights into plant adaptation strategies under increasing environmental stress.

## Materials and Methods

### Plant Material, Seed Sowing, and Germination

Rapeseed (*Brassica napus* L.) spring variety Helios seeds were obtained from the Germplasm Resource Information Network (GRIN, Czech Republic). Two seeds were sown per pot (11 × 11 × 18 cm) filled with a 1:1 mixture of peat and perlite, moistened with Hoagland solution (Malhotra et al., 2014). The pots were placed in a growth room under a 16-hour light / 8-hour dark cycle at 22 °C, 50–55% relative humidity, and 150 μmol m^-2^ s^-1^ light intensity. Pots were watered every three days with either Hoagland solution or distilled water.

## Cold Stress Treatment

The experimental design is illustrated in Fig. 1. Previous studies confirmed that Helios exhibits cold hardiness and is vernalization responsive (Çağlı et al., 2025). At five to six weeks after sowing, when seedlings reached the four-to five-leaf stage (BBCH scale 14–15), twenty pots were transferred to a growth chamber (Nüve TK 252) set at 4 °C under short-day conditions for three weeks. After cold exposure, these plants were returned to the growth room at 22 °C. Another twenty pots remained in the growth room at 22 °C without cold treatment.

**Fig. 1.**
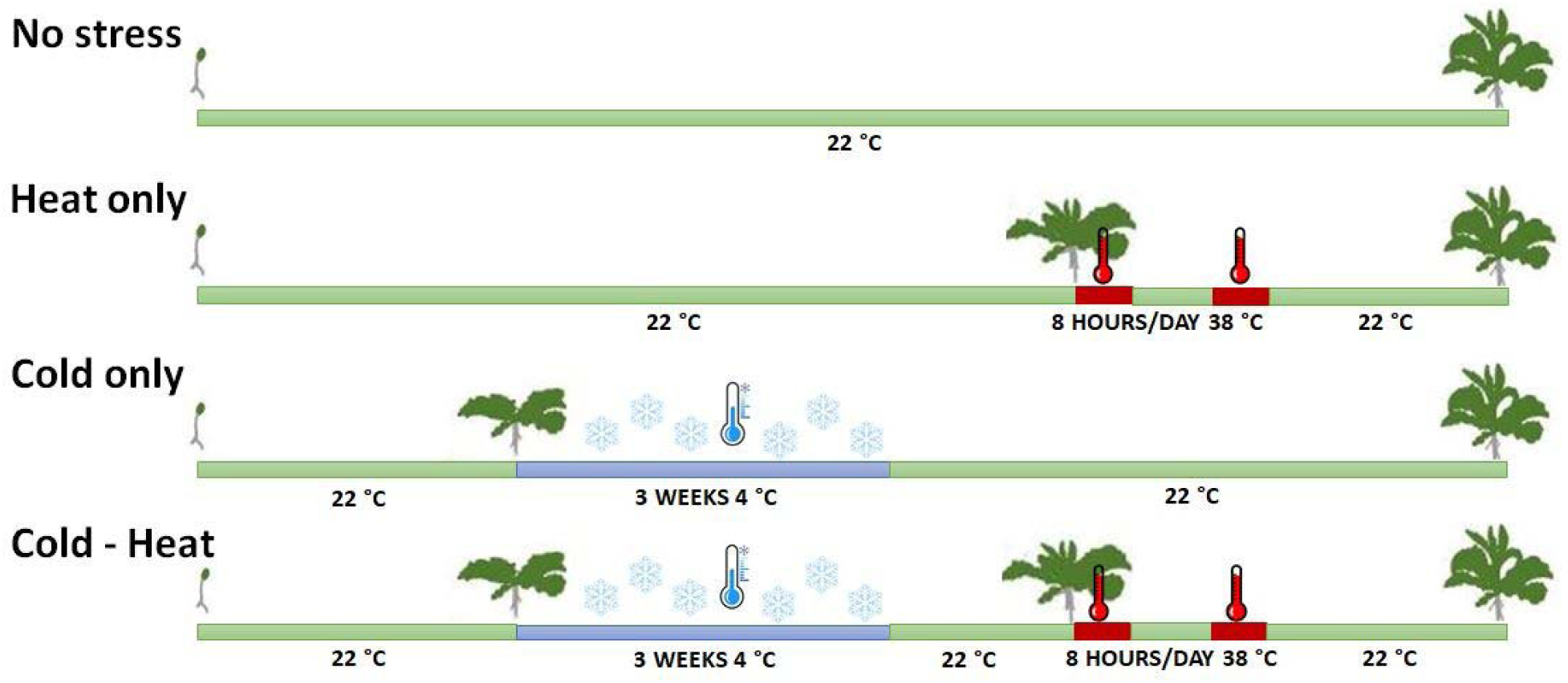
The experimental set-up. Under ‘No Stress’, plants were maintained at 22 °C throughout the entire growth period. ‘Heat only’ plants were grown at 22°C until they reached the 8-9 leaf stage, then exposed to 38° C for 8 hours per day over two consecutive days and returned to 22 °C. ‘Cold only’ plants were grown at 22°C until the 4-5 leaf stage, followed by three weeks-cold period at 4 °C. Under the ‘Cold-Heat’ treatment, after the three weeks-cold stress, plants were returned to the growth room at 22°C until they reached 8-9 leaf stage and subjected to heat stress at 38 °C for 8 hours per day over two consecutive days.

### Heat Stress Treatment

At the eight- to nine-leaf stage (BBCH scale 18–19), ten pots previously exposed to cold stress and ten non-cold-treated pots were subjected to heat stress in a growth chamber at 38 °C for 8 hours daily over two consecutive days, under light and 70% relative humidity. Following heat stress, pots were returned to the growth room at 22 °C. Twenty pots without heat stress (with or without prior cold treatment) remained in the growth room.

### Growth Conditions to Full Maturity

On January 19, 2023, eight days after heat stress ended, all forty pots representing the four treatment groups (‘No stress’, ‘Cold only’, ‘Heat only’, and ‘Cold – Heat’) were transferred to a semi-controlled greenhouse equipped with indoor heating and supplemented with 150 W LED lighting (Philips Xitanium) as needed. At transplantation, plants were at the nine- to ten-leaf stage. Flowering time was recorded as the number of days from transplantation until the first flower appeared. Physiological parameters—plant height (cm), number of lateral branches, number of pods on the main stem, number of seeds per pod, and thousand-seed weight (g)—were measured. Mean values were calculated from at least three biological replicates for each treatment group.

### Seed Germination Assay

One hundred seeds from each treatment group were surface sterilized in 5% sodium hypochlorite (v/v) and 2.5% Tween 20 (v/v) solution for 15 minutes with constant shaking at room temperature. Seeds were rinsed five times with sterile distilled water and dried. Fifty seeds per group were placed on glass petri dishes lined with double filter paper moistened with half-strength Hoagland’s solution. Seeds were stratified at 4 °C for four days and then incubated in a growth chamber at 23 °C and 60% relative humidity for 10 days. Germinated seedlings were counted and photographed. Seed germination percentage was calculated as:

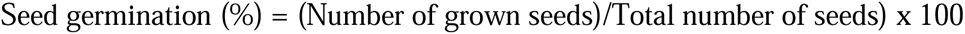

### Determination of Total Phenolic and Flavonoid Contents

For seed extracts, 100 mg of seeds were ground and mixed with 1 ml of 80% methanol. Samples were sonicated for 45 minutes and centrifuged at 10,000 g for 30 minutes. The supernatant was mixed 1:1 with 80% methanol (Eynck et al., 2009). Total phenolic content (TPC) was measured by mixing 30 µl of extract with Folin-Ciocalteu reagent and other reagents, incubating as described, and measuring absorbance at 725 nm against a gallic acid standard curve.

Total flavonoid content (TFC) was measured by reacting 100 µl of extract with 100 µl of 2% AlCl_3_ solution, incubating in the dark for 15 minutes, and measuring absorbance at 430 nm (Elfalleh et al., 2009). Quercetin was used for the calibration curve.

### DPPH Radical Scavenging Assay

Free radical scavenging activity was assessed by the DPPH assay (Prieto, 2012). Extract stock solutions (50 mg/mL) were serially diluted. In 96-well plates, 100 µL of each dilution was mixed with 100 µL of 0.2 mM DPPH in methanol. After 30 min incubation in the dark at room temperature, absorbance was measured at 517 nm. Scavenging activity (%) was calculated as:

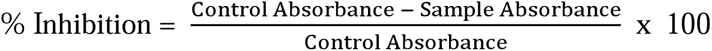

IC_50_ values (concentration scavenging 50% of DPPH radicals) were determined from dose-response curves.

### Ferric Reducing Antioxidant Power (FRAP) Assay

FRAP assay was performed according to Xiao et al. (2020). Six microliters of seed extract were mixed with 200 µl FRAP working solution and incubated at 37 °C in the dark for 30 minutes. Absorbance was measured at 593 nm. A calibration curve was prepared with FeSO_4_·7H_2_O (0–4 mM). Results were expressed as µmol Fe^2+^ equivalents per gram of dry weight (DW) of the extract. Samples were analyzed in triplicate.

### Hydrogen Peroxide Content

H_2_O_2_ content was determined per Velikova et al. (2000). Frozen seed powder (100 mg) was mixed with 5 ml 0.1% trichloroacetic acid, centrifuged, and the supernatant reacted with 1 mL of 1M KI, and 0.5 mL of 10 mM phosphate buffer (pH 7.0). After 60 minutes’ incubation in the dark, absorbance was read at 390 nm. H_2_O_2_ concentration was calculated against a standard curve.

### Superoxide Dismutase (SOD) and Catalase (CAT) Activity Assays

Seed extracts were prepared by grinding 100 mg seeds with sodium phosphate buffer containing polyvinylpolypyrrolidone (Önder et al., 2020). The homogenate was centrifuged at 27,000 g for 50 minutes at 4 °C, and the supernatant was collected. Protein concentration was measured by Qubit fluorometer (Thermo Fisher Scientific, USA). SOD activity was measured in a reaction mixture with 13 mM methionine, 10 µM EDTA, 50 mM sodium phosphate (pH 7.8), 2 µM riboflavin, 75 µM of nitro blue tetrazolium and 100 µl of enzyme extract after incubation under light for 10 minutes. The absorbance was measured at 560 nm. SOD activity was expressed as U mg^-1^ protein.

CAT activity was assayed by monitoring decomposition of H_2_O_2_ at 240 nm over one minute in a reaction buffer containing 0.5 mL of 40 mM H_2_O_2_, 2 mL sodium phosphate buffer (pH 7.0), and 0.5 mL enzyme extract (Önder et al., 2020). One unit of enzyme activity equals µmol H_2_O_2_ decomposed.

### Chlorophyll Content

Chlorophyll *a*, *b*, and total chlorophyll were measured following Arnon (1949). Fresh leaves (0.25 g) of rapeseed seedlings-germinated from seeds of four treatment groups-were ground in liquid nitrogen and extracted with 80% acetone. Absorbance at 645 and 663 nm was measured. Chlorophyll contents (mg g^-1^) were calculated using:

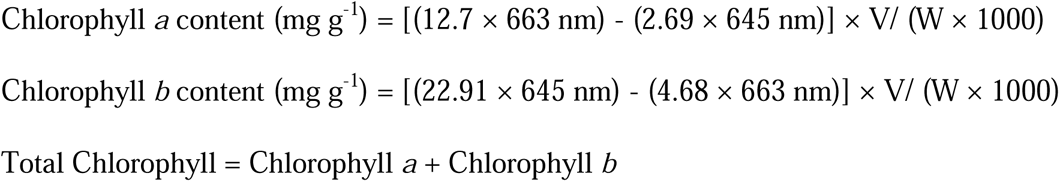

Where V = volume of extract (mL), W = fresh weight of tissue (μg).

### Relative Water Content (RWC)

True leaves of seedlings were excised and the fresh weight (FW) was recorded. Samples were soaked in distilled water under light for 24 hours to measure turgor weight (TW), then oven-dried at 80 °C for 48 hours to determine dry weight (DW). Relative water content (%) was calculated using the following formula (Ritchie et al., 1990):

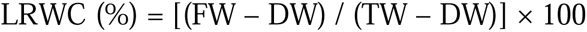

### Fresh Seedling Weight

Nine seeds per biological replicate were sown in trays (3 seeds/tray), kept in the dark at 20 °C until germination, then under light at the same temperature for 20 days. Seedlings were watered every two days, harvested carefully, washed, dried, and fresh weights measured.

### Total Lipid Extraction and Oil Content

Total lipid extraction followed Çağlı et al. (2025). Briefly, mature seeds were oven-dried (42 °C, 150 min), placed in a desiccator for 48 h, and weighed. Ten mg seed samples were mixed with 2 ml isopropanol, heated at 85 °C for 10 minutes, homogenized, then 3 ml hexane was added and vortexed. After incubation and sodium sulfate addition, the upper phase was collected and evaporated. Oil content (%) was determined gravimetrically.

### Fatty Acid Methyl Ester (FAME) Analysis

FAME compositions of seeds from ‘no stress’, ‘cold only’, ‘heat only’ and cold – heat’ treatments were analyzed by gas chromatography (GC) as described in Çağlı et al. (2025). Lipid extracts were transesterified with 2 N potassium hydroxide, mixed with hexane, centrifuged at 3,000 g for 15 minutes. The upper phase was analyzed using a SHIMADZU GCMS-QP2020 with RESTEK Rxi-5 Sil MS column at 250 °C and 34.17 psi. FAMEs were quantified as percentage peak areas compared to standards.

### RNA Extraction and qRT-PCR

Total RNA was extracted from developing seeds collected at BBCH scale 80 following Çağlı et al. (2025). Samples were homogenized in 1 mL RNA extraction buffer, treated with acidic phenol:chloroform (pH 4.3), precipitated with isopropanol and sodium acetate, washed, and dissolved in DEPC-treated water. RNA was DNase-treated (Turbo DNA-free kit). cDNA synthesis used 1 µg RNA with iScript cDNA kit (Bio-Rad). qRT-PCR reactions (15 µL) contained SYBR Green mix and 6 µL diluted cDNA. Expression was normalized to Ubiquitin using the comparative Ct method. Primers for *BnaFAD2*, *BnaFAD5*, *BnaFATB*, *BnaMCOA* (*AAE13*), *BnaWD40*, and Ubiquitin were as listed in Çağlı et al. (2025).

### Statistical Analysis

Experiments included at least three biological replicates. Data are presented as mean ± standard error (SE). One-way ANOVA and unpaired Mann-Whitney U tests were used for comparisons. Statistical significance was set at *p < 0.05*. Pearson’s correlation coefficient (*r*) was calculated for parameter correlations. Analyses were performed using GraphPad Prism 9.

## Results

### Experimental Design of Stress Treatments

Four temperature stress treatments were applied to *Brassica napus* cv. Helios plants: optimal control (22 °C), heat stress (38 °C, 8 h/day for 2 days), cold stress (4 °C for 3 weeks), and a sequential cold–heat regime (3 weeks cold followed by heat stress) (Fig. 1). This design aimed to evaluate physiological, biochemical, and molecular responses in the progeny of stressed plants and their first stress-free generation, testing for potential priming effects.

### Physiological Responses to Temperature Stress

Temperature stress significantly influenced key agronomic traits (Table 1). Cold stress shortened flowering time (FT) (57.2 ± 3.3 days vs. 71.7 ± 1.9 days in control, *p < 0.01*), while ‘heat only’ (73.7 ± 1.9 days) and cold–heat (65.6 ± 2.2 days) treatments delayed or moderated flowering, respectively. The difference in FT between ‘cold only’ and ‘cold – heat’ groups was statistically significant (*p < 0.05*). Plant height showed slight, non-significant increases under heat (124.7 ± 4.1 cm) and cold (123.0 ± 2.3 cm) stresses compared to ‘no stress’ (118.3 ± 2.0 cm). Lateral branching decreased significantly under stress, especially cold (7.0 ± 0.6 vs. 10.0 ± 0.0 in control; *p < 0.05*). Thousand seed weight dropped across all stresses, most notably in the cold–heat group (0.16 ± 0.01 g vs. 0.28 ± 0.01 g in control; *p < 0.01*). Number of pods on the main stem increased under heat but decreased under cold–heat treatment (565.0 ± 29.9 vs. 442.3 ± 6.6, respectively; *p < 0.05*) relative to control (485.7 ± 19.4). Seed number per pod remained unchanged.

**Table 1.** Physiological parameters (flowering time (FT), plant height (PH), lateral branches (LB), Thousand Seed Weight (TSW), number of pods on the main stem (NOPOTMS), and seed number per pod (SNPP)) of no stress, heat only, cold only, and cold - heat stress treatment groups.

|  | **FT** | **PH** | **LB** | **TSW** | **NOPOTMS** | **SNPP** |
| --- | --- | --- | --- | --- | --- | --- |
| **No stress** | 71.71 ± 1.87<sup>a</sup> | 118.33 ± 2.03 | 10.00 ± 0.00<sup>c</sup> | 0.28 ± 0.01<sup>d</sup> | 485.67 ± 19.35 | 22.73 ± 3.82 |
| **Heat only** | 73.67 ± 1.89 | 124.67 ± 4.07 | 08.00 ± 0.58 | 0.18 ± 0.01<sup>d</sup> | 565.00 ± 29.86<sup>f</sup> | 23.47 ± 1.90 |
| **Cold only** | 57.22 ± 3.25<sup>a, b</sup> | 123.00 ± 2.31 | 07.00 ± 0.58<sup>c</sup> | 0.23 ± 0.19<sup>e</sup> | 473.33 ± 08.74 | 23.13 ± 5.80 |
| **Cold - Heat** | 65.58 ± 2.19<sup>b</sup> | 119.00 ± 1.73 | 09.00 ± 0.58 | 0.16 ± 0.01<sup>f</sup> | 442.33 ± 06.61<sup>f</sup> | 25.20 ± 4.66 |

Values are represented as mean ± SEM. b, c, e, f: $p < 0.05$ ; a, d: $p < 0.01$

### Total Phenolic and Flavonoid Contents

Temperature stress during the vegetative growth phase of rapeseed plants significantly altered the total phenolic (TPC) and total flavonoid contents (TFC) in seeds (Fig. 2). Heat stress markedly increased TPC (4.67 ± 0.08 GAE/g DW) and TFC (0.44 ± 0.01 mg QUE/g DW) compared to control (TPC: 3.98 ± 0.11 mg GAE/g DW, *p < 0.001*; TFC: 0.29 ± 0.01 mg QUE/g DW, *p < 0.001*). Among all treatments, the lowest TPC was recorded in the seeds of ‘cold only’ plants (3.66 ± 0.03 mg GAE/g DW) and TFC was similar to ‘no stress’ levels (0.28 ± 0.02 mg QUE/g DW). Sequential cold–heat treatments produced intermediate levels of TPC (4.29 ± 0.10 mg GAE/g DW; *p < 0.05*) and TFC (0.37 ± 0.01 mg QUE/g DW; *p < 0.05*), indicating partial mitigation of cold’s suppressive effect by subsequent heat.

**Fig. 2.**
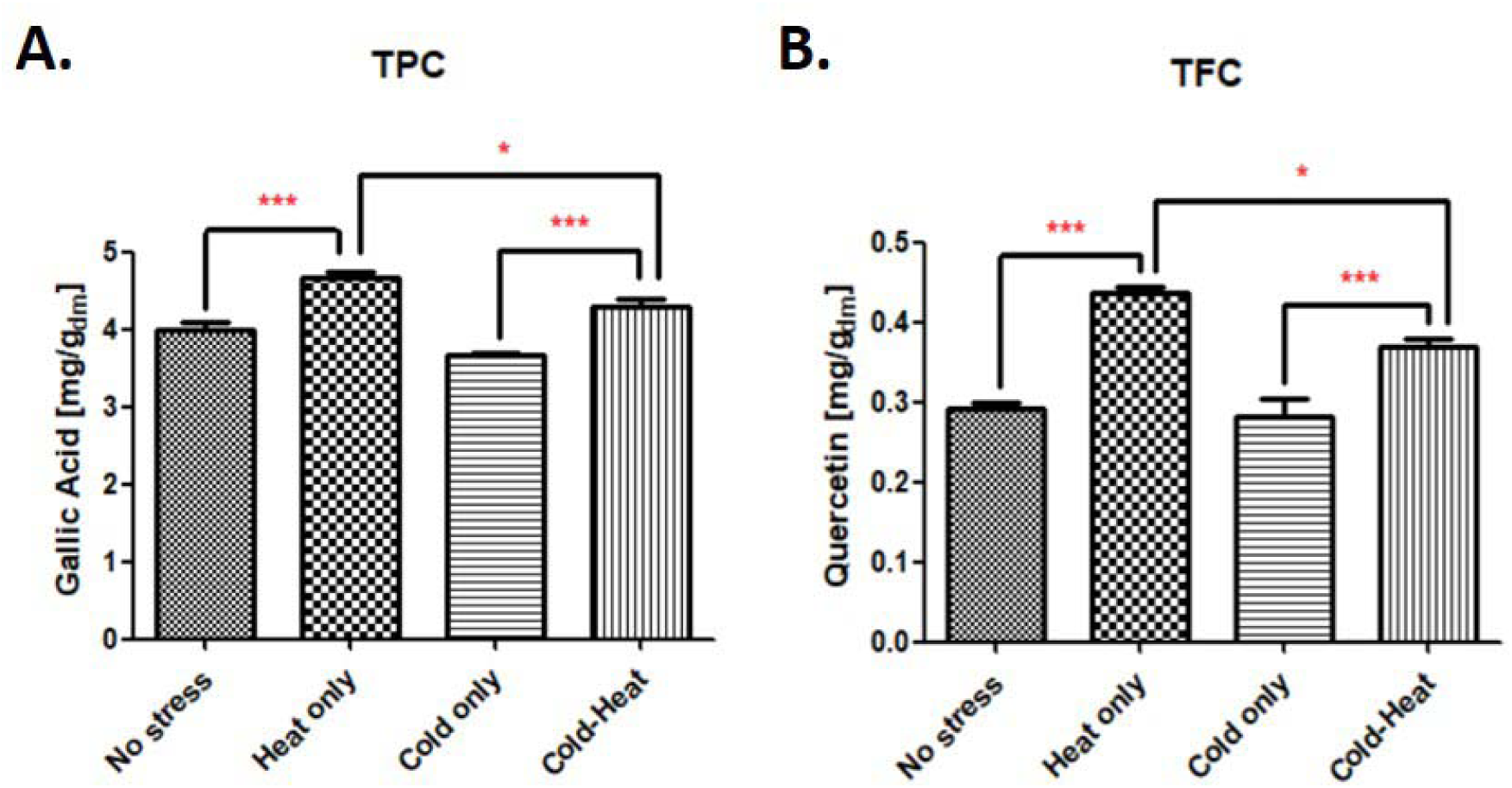
Total phenolic (TPC) and total flavonoid content (TFC) of *B. napus* seeds that were exposed to no stress, heat stress only, cold stress only and cold - heat stresses. **(A)** TPC is expressed as mg gallic acid equivalents (GAE) per gram of dry weight; **(B)** TFC is expressed as mg quercetin equivalents (QUE) per gram of dry weight. Values are represented as mean ± SEM. * *p < 0.05*, ** *p < 0.01*, *** *p < 0.001*

### Antioxidant Activities

Antioxidant capacity, measured by DPPH and FRAP assays, increased under heat and cold–heat stress (Fig. 3). Heat-treated seeds showed lower DPPH IC_50_ values (1.37 ± 0.07 mg/g DW vs. 2.04 ± 0.11 control*, p < 0.001*) and higher FRAP values (16.87 ± 0.15 mg FeSO_4_/g DW vs. 14.86 ± 0.24 control, *p < 0.001*). A similar reduction in IC_50_ values was observed in the cold–heat group (1.26 ± 0.10 mg/g DW), and ferric reducing antioxidant power (17.83 ± 0.23 mg/g FeSO_4_ DW; *p < 0.001*) was elevated. Cold stress alone did not significantly enhance antioxidant activity. These results indicate that heat stress alone or following cold stress elevates the antioxidant activities of *B. napus* seeds.

**Fig. 3.**
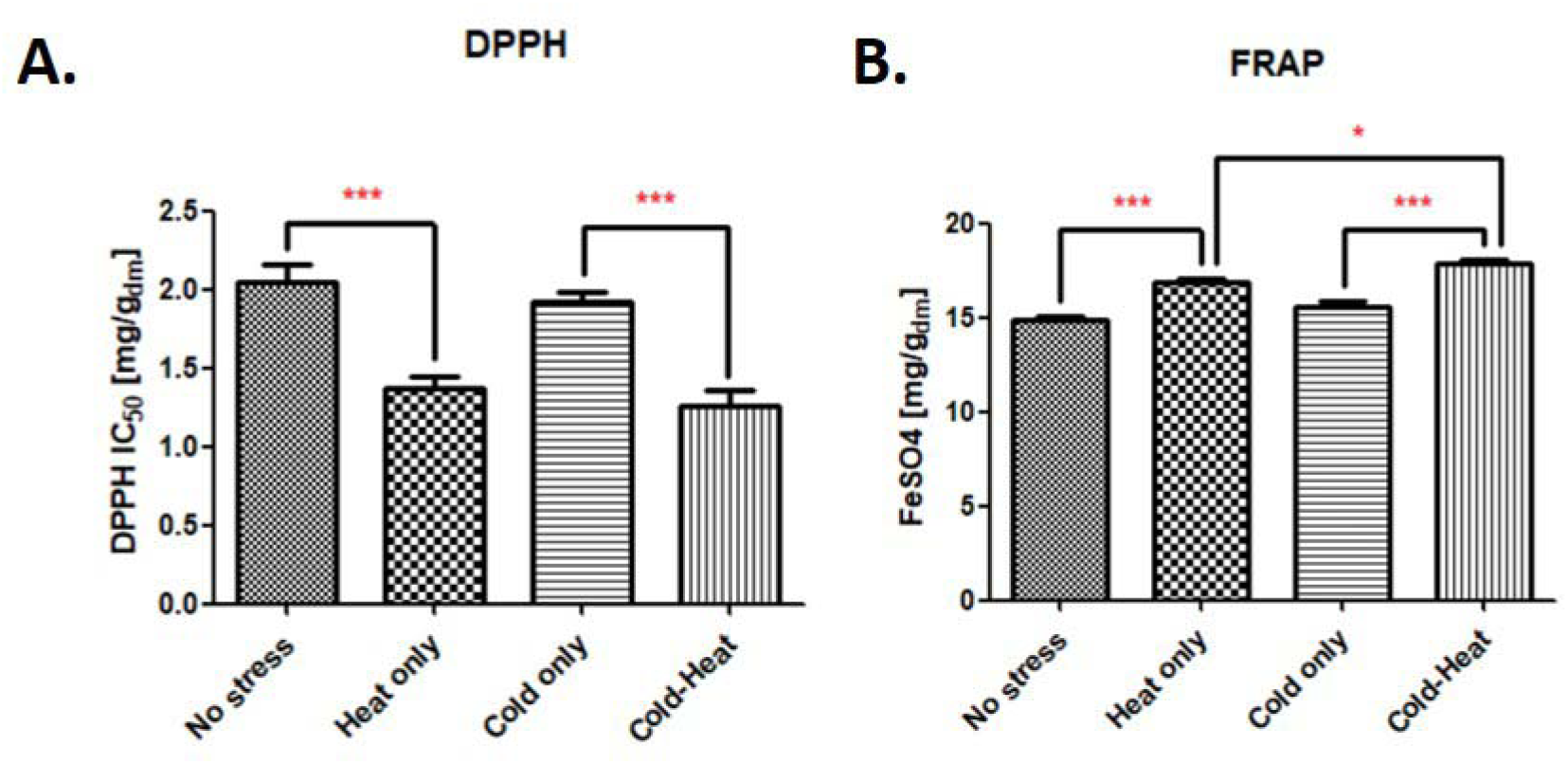
Antioxidant activities of *B. napus* seeds that were exposed to no stress, heat stress only, cold stress only and cold - heat stresses **(A)** 2,2-Diphenyl-1-picrylhydrazyl (DPPH) radical scavenging activity is expressed as the IC_50_ value (µg/mL) of the plant dry weight, **(B)** Ferric reducing antioxidant power (FRAP) is expressed as mg FeSO_4_ per gram of dry weight. Values are represented as mean ± SEM. * *p < 0.05*, ** *p < 0.01*, *** *p < 0.001*

### Correlations Between Phenolics, Flavonoids, and Antioxidant Activities

We calculated the Pearson correlation coefficients (*r*) to assess the correlations between TPC and TFC, DPPH and FRAP assay results (Fig. 4). TPC and TFC were strongly positively correlated (*r* = 0.97, *p < 0.05*). Both negatively correlated with DPPH IC_50_ (indicative of increased antioxidant capacity), though not significantly. A strong negative correlation existed between DPPH IC_50_ and FRAP (*r* = –0.98, *p < 0.05*), reflecting consistent antioxidant assay results. Moderate positive correlations of TPC and TFC with FRAP were observed but were not significant. These findings suggest that in addition to TPC and TFC, other factors may also contribute to the antioxidant acitivity of *B. napus* seeds.

**Fig. 4.**
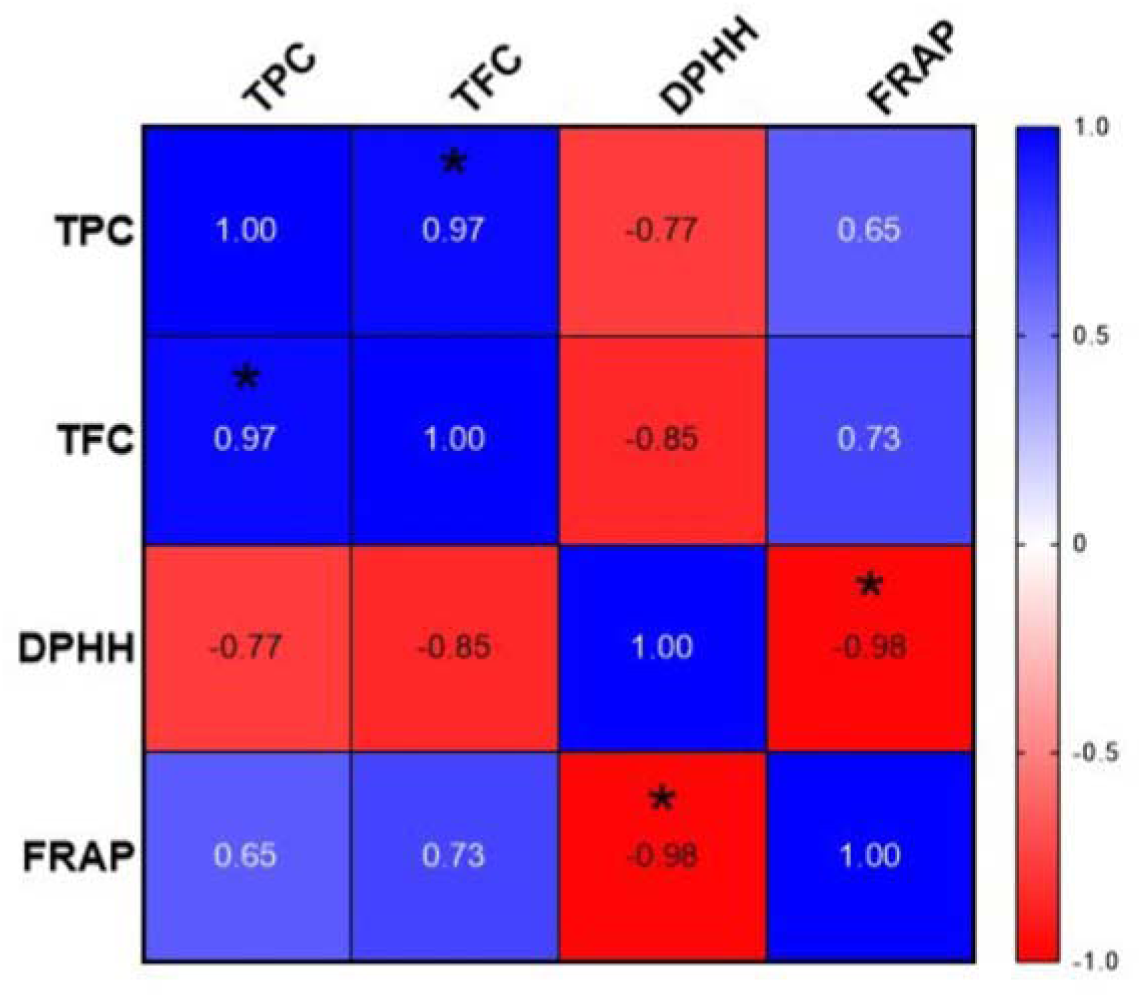
Pearson correlation coefficients (*r*) between four variables; Total Phenolic Content (TPC), Total Flavonoid Content (TFC), 2,2-Diphenyl-1-picrylhydrazyl (DPPH), and Ferric Reducing Antioxidant Power (FRAP). * *p < 0.05*

### Enzymatic Antioxidant Activities and H_2_O_2_ Content

Superoxide dismutase (SOD) activity exhibited a significant increase across all stress treatments compared to ‘no stress’ control (27.32 ± 0.44 U mg^-1^ protein) (Table 2). The highest SOD activity was observed in the cold–heat group (45.03 ± 1.82 U mg^-1^ protein; *p < 0.05*), followed by ‘heat only’ (36.95 ± 1.21 U mg^-1^ protein) and ‘cold only’ (35.58 ± 2.57 U mg^-1^ protein) treatments. Catalase (CAT) activity was slightly lower across all treatments compared to control (0.021 ± 0.002 U mg^-1^ protein). Hydrogen peroxide levels remained stable across treatments showing no significant changes compared to control (8.20 ± 0.05 µmol g^-1^).

**Table 2.** Superoxide dismutase (SOD) and catalase (CAT) activities and H_2_O_2_ content of seeds under four treatment conditions: No stress, Heat only, Cold only, and Cold - Heat.

|  | **SOD Activity**<br>(U mg<sup>-1</sup> protein) | **CAT Activity**<br>(U mg<sup>-1</sup> protein) | **H<sub>2</sub>O<sub>2</sub> Content**<br>(μmol g<sup>-1</sup>) |
| --- | --- | --- | --- |
| **No stress** | 27.32 ± 0.44<sup>a-b</sup> | 0.021 ± 0.002 | 8.20 ± 0.05 |
| **Heat only** | 36.95 ± 1.206<sup>a</sup> | 0.016 ± 0.001 | 7.97 ± 0.38 |
| **Cold only** | 35.58 ± 2.569<sup>b-c</sup> | 0.014 ± 0.001 | 8.42 ± 0.05 |
| **Cold -Heat** | 45.03 ± 1.819<sup>c</sup> | 0.018 ± 0.002 | 7.79 ± 0.11 |

Data are represented as mean ± standard error of the mean (SEM). a, b, c: $p < 0.05$

### Seed Oil Content and Fatty Acid Composition

Cold stress significantly increased total seed oil content (43.61 ± 1.82% vs. 35.56 ± 0.57% control, *p < 0.05*), while heat and cold–heat treatments showed slight or no increases (Table 3). Oleic acid decreased under all stresses, especially cold (60.99 ± 0.46% vs. 66.49 ± 1.48% control), whereas linoleic acid and total polyunsaturated fatty acids (PUFAs) increased with cold stress, suggesting enhanced membrane fluidity and stress tolerance (Tables 3 and S1).

**Table 3.** Seed Oil Content (% OIL), Oleic Acid Content (% OA), and Linoleic Acid Content (% LA) of No stress, Heat only, Cold only, Cold - Heat stress treatment groups.

|  | **% OIL** | **% OA** | **% LA** |
| --- | --- | --- | --- |
| **No stress** | 35.56 ± 0.57<sup>a</sup> | 66.49 ± 1.48 | 18.91 ± 1.02 |
| **Heat only** | 38.28 ± 0.77 | 64.55 ± 1.63 | 20.19 ± 1.22 |
| **Cold only** | 43.61 ± 1.82<sup>a-b</sup> | 60.99 ± 0.46 | 22.57 ± 0.47 |
| **Cold - Heat** | 35.00 ± 2.25<sup>b</sup> | 63.00 ± 0.17 | 19.77 ± 0.11 |

Data are represented as mean ± standard error of the mean (SEM). a, b: $p < 0.05$

### Expression of Fatty Acid Synthesis Genes

Early temperature stress significantly affected the expression of fatty acid synthesis-related genes—*BnaFAD2*, *BnaFAD5*, *BnaFATB*, *BnaMCOA*, and *BnaWD40*—in developing seeds (Fig. 5). Sequential cold–heat treatment maintained higher *BnaFAD2* levels than heat stress alone (*p < 0.05*) (Fig. 5A). *BnaFAD5* was strongly induced by cold stress (*p < 0.001*) and moderately by heat stress (*p < 0.05*) (Fig. 5B). Similarly, *BnaFATB* showed marked upregulation in response to cold stress (*p < 0.01*), but expression returned to near-control levels under combined treatment (Fig. 5C). *BnaMCOA* and *BnaWD40* exhibited moderate changes across treatments; however, *BnaWD40* was significantly upregulated in the cold–heat condition (*p < 0.05*) (Fig. 5D and E).

**Fig. 5.**
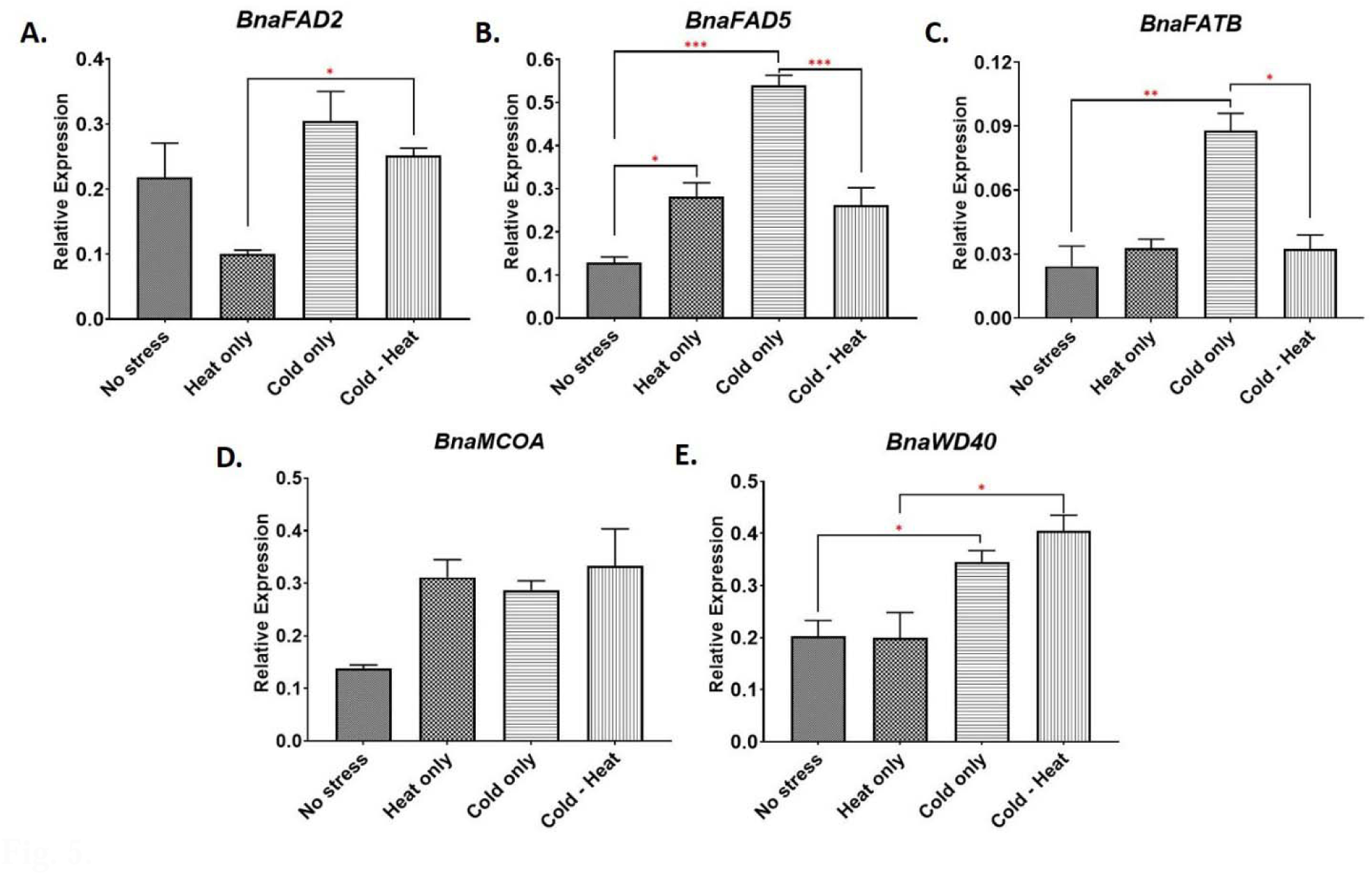
Expression profiles of fatty acid synthesis-related genes, **(A)** *BnaFAD2*, **(B)** *BnaFAD5*, **(C)** *BnaFATB*, **(D)** *BnaMCOA*, and **(E)** *BnaWD40* are shown as bar graphs of mean values ± SEM for three biological replicates. Statistical significance was determined based on p-values: * *p < 0.05*, ** *p < 0.01*, *** *p < 0.001*

These findings suggest that temperature stress differentially regulates key genes in fatty acid metabolism, with cold stress serving as a more potent inducer than heat. The results also support the possibility of transcriptional memory persisting through both mitosis and meiosis in response to thermal stress.

### Correlation Between Gene Expression and Oil Composition

Pearson correlation coefficients (*r*) were calculated to assess relationships between the expression of fatty acid synthesis-related genes, total seed oil content, and oleic acid content in *B. napus* under temperature stress (Fig. 6). *BnaFAD5* expression showed a strong positive correlation with *BnaFATB* (*r =* 0.97, *p < 0.05*) and a strong negative correlation with oleic acid content (*r* = - 0.99, *p < 0.05*). It also exhibited a positive, though not statistically significant, correlation with total oil content (*r* = 0.92). Similarly, *BnaFATB* was positively correlated with total oil content (*r =* 0.94) and negatively with oleic acid content (*r* = −0.94), but these associations were not significant. *BnaMCOA* and *BnaWD40* showed moderate positive correlations with other genes, yet no significant links to oil or oleic acid levels. Notably, total oil content was strongly and negatively correlated with oleic acid percentage (*r* = −0.85), suggesting that stress-induced oil accumulation may be accompanied by a shift toward polyunsaturated fatty acids. These results underscore *BnaFAD5* as a potential regulatory gene influencing both oil quantity and fatty acid composition under temperature stress.

**Fig. 6.**
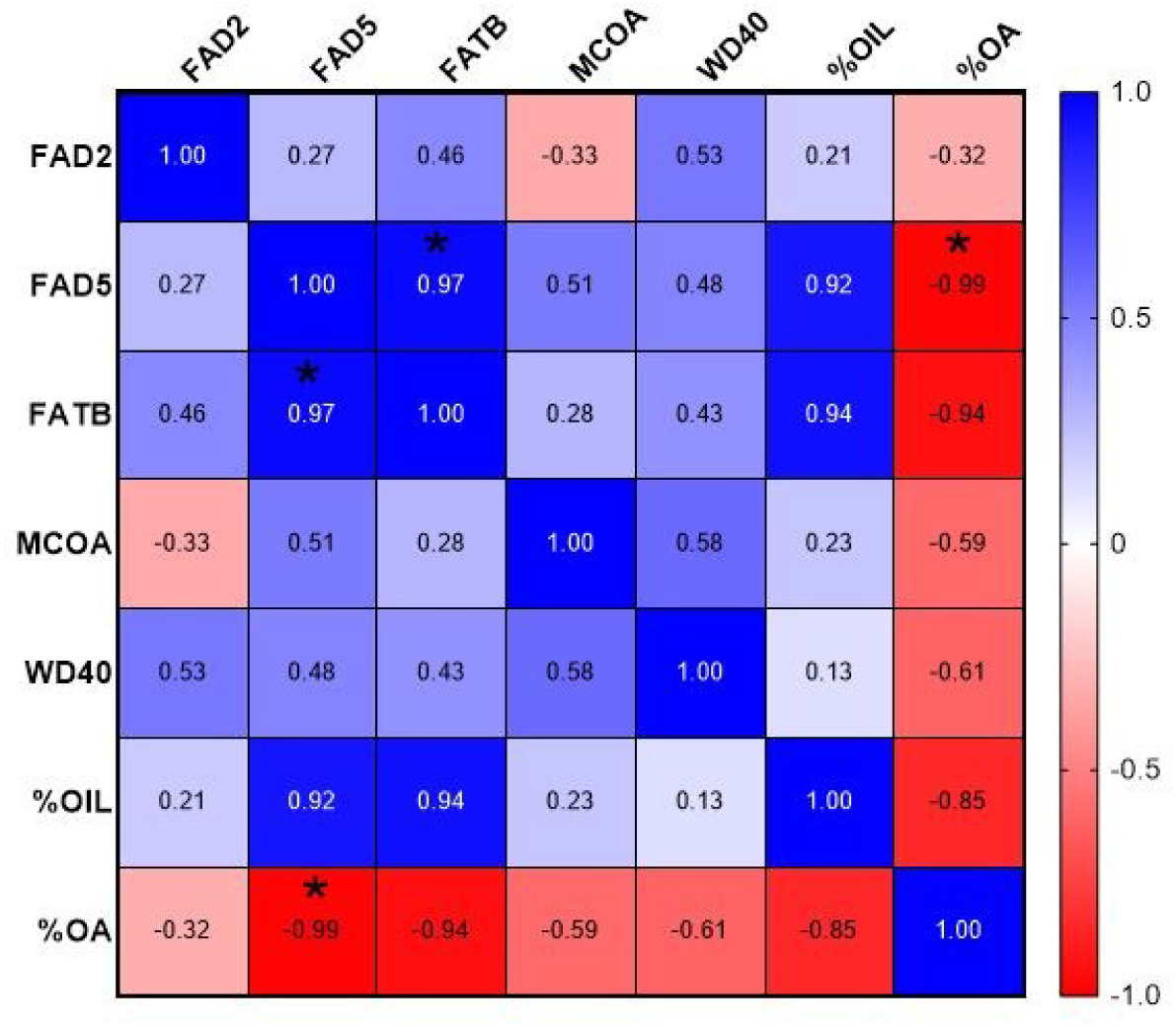
Pearson correlation coefficients (*r*) between fatty acid-related gene expressions, seed oil content, and oleic acid content. * *p < 0.05*

### Seed Germination

Germination analysis of seed progeny from stress-exposed plants showed minimal variation across treatments, with all groups exhibiting high germination rates (Table 4). Control seeds germinated at 97.667 ± 0.002%, while seeds from heat-stressed plants showed a similar rate of 97.334 ± 0.003%. Cold stress slightly reduced germination to 95.667 ± 0.002%, and the combined cold–heat treatment yielded 95.334 ± 0.001%. Despite these minor differences, all treatments maintained germination above 95%, indicating that temperature stress during seed development did not significantly affect seed viability.

**Table 4.** Analysis of average germination percentage of rapeseed under different treatment conditions.

|  | **Germination percentage (G%)** |
| --- | --- |
| **No stress** | 97.667 ± 0.002 |
| **Heat only** | 97.334 ± 0.003 |
| **Cold only** | 95.667 ± 0.002 |
| **Cold - Heat** | 95.334 ± 0.001 |

Values are means of three biological and two technical replicates. Data are represented as Mean ± SEM.

### Chlorophyll Content in Stress-Free Progeny

Chlorophyll measurements in the first stress-free generation of rapeseed seedlings revealed significant differences among treatment groups (Table 5). Total chlorophyll content was highest in progeny of plants exposed to combined cold–heat stress (1.063 ± 0.00001 µg g^-1^; *p < 0.01*) compared to controls (0.902 ± 0.00001 µg g^-1^). Heat-stressed progeny showed a slight increase (0.922 ± 0.00001 µg g^-1^), while cold stress alone resulted in the lowest total chlorophyll content (0.864 ± 0.00004 µg g^-1^; *p < 0.001*). This pattern was mirrored in chlorophyll *a* and *b* levels. Chlorophyll *a* increased in the heat-only (0.620 ± 0.00001 µg g^-1^) and cold–heat groups (0.675 ± 0.00002 µg g^-1^; *p < 0.01*) but decreased in cold-only progeny (0.566 ± 0.00002 µg g^-1^) relative to controls (0.589 ± 0.00002 µg g^-1^). Chlorophyll *b* peaked in cold–heat plants (0.398 ± 0.00001 µg g^-1^; *p < 0.001*), while heat-only (0.301 ± 0.00000 µg g^-1^) and cold-only (0.298 ± 0.00002 µg g^-1^) treatments showed reductions compared to control (0.317 ± 0.00000 µg g^-1^). These results suggest that sequential cold–heat stress may prime progeny for enhanced chlorophyll accumulation, potentially improving photosynthetic capacity.

**Table 5.** Chlorophyll *a*, Chlorophyll *b*, and Total Chlorophyll Contents (µg.g^-1^) of first stress-free generation of rapeseed seedlings under different treatment conditions.

|  | **Chlorophyll *a***<br>**content ($\mu\text{g g}^{-1}$)** | **Chlorophyll *b* content**<br>**($\mu\text{g g}^{-1}$)** | **Total Chlorophyll**<br>**Content ($\mu\text{g 1g}^{-1}$)** |
| --- | --- | --- | --- |
| **No stress** | $0.589 \pm 0.00002^{\text{a}}$ | $0.317 \pm 0.00000^{\text{c}}$ | $0.902 \pm 0.00001^{\text{e}}$ |
| **Heat only** | $0.620 \pm 0.00001$ | $0.301 \pm 0.00000$ | $0.922 \pm 0.00001$ |
| **Cold only** | $0.566 \pm 0.00002^{\text{b}}$ | $0.298 \pm 0.00002^{\text{d}}$ | $0.864 \pm 0.00004^{\text{f}}$ |
| **Cold - Heat** | $0.675 \pm 0.00002^{\text{a-b}}$ | $0.398 \pm 0.00001^{\text{c-d}}$ | $1.063 \pm 0.00001^{\text{e-f}}$ |

Values are represented as Mean ± SEM. a: $p < 0.05$ ; b, c, e: $p < 0.01$ ; d, f: $p < 0.001$

### Fresh Seedling Weight and Leaf Relative Water Content

Fresh weight measurements of the first stress-free generation seedlings showed minor differences among treatment groups (Table S2). Progeny of plants exposed to combined cold–heat stress had the highest total fresh seedling weight (3.765 ± 0.015 g) and average weight per seedling (0.418 ± 0.002 g), compared to control (3.180 ± 0.080 g total; 0.353 ± 0.009 g per seedling). Heat and cold stress alone also led to slight increases in fresh weight relative to control.

Leaf relative water content (LRWC) varied among treatment groups in the first stress-free generation of *B. napus* seedlings (Table S3). Progeny of combined cold–heat stressed plants had the highest LRWC (90.37 ± 0.78%), followed by cold-only plants (85.22 ± 1.47%). Control seedlings showed 80.27 ± 0.97%, while heat-only progeny had similar values (79.82 ± 3.17%). Taken together, combined stress exposure may induce intergenerational effects that enhance biomass accumulation and relative water content in the progeny.

## Discussion

The capacity of plants to develop cross-tolerance to multiple abiotic and biotic stresses represents a vital adaptive strategy, enhancing resilience and productivity in fluctuating environments. In this study, we investigated the effects of single and sequential temperature stresses—cold, heat, and combined cold–heat—applied during the vegetative phase of *Brassica napus* (cv. Helios), with a focus on their impact on seed quality and the intergenerational consequences.

Although Helios has previously been characterized as a spring variety (Jørgensen & Andersen, 1994; Brown et al., 1995), our findings confirm that the seedlings display winter hardiness and vernalization responsiveness (Çağlı et al., 2025). Three-week cold exposure significantly accelerated flowering in Helios plants through cold-induced epigenetic silencing of the major floral repressor, *FLC* orthologs in rapeseed (De Lucia et al., 2008; Menon et al., 2021; Çağlı et al., 2025). Interestingly, sequential cold–heat exposure reversed this acceleration, delaying flowering relative to cold alone. This observation aligns with reports in *Brassica rapa* where elevated temperatures suppress *flowering locus T* (*FT*) expression via increased H2A.Z deposition to delay flowering (Del Olmo et al., 2019). In *B. napus*, elevated temperatures also delay flowering time, however; the involvement of H2A.Z dynamics at the *FT* locus is genotype-dependent (Abelenda et al., 2023). These findings highlight a potentially complex interaction between vernalization pathways and heat stress responses, possibly mediated by chromatin dynamics, which requires further investigation.

Seed yield is another important agronomic trait that is highly dependent on the reproductive success of plants. Sequential cold–heat stress negatively affected yield parameters, including thousand seed weight and number of pods on the main stem. While the independent effects of heat or cold on reproductive traits have been well documented, our data show that their combination further exacerbates reproductive stress, leading to reduced seed production (Yu et al., 2014; Huang et al., 2019; Chen et al., 2021; Chen et al., 2024).

ROS and antioxidant defenses play a central role in cross-tolerance and transgenerational memory (Mei et al., 2010; Li et al., 2014; van Buer et al., 2016; Hossain et al., 2018; Liu et al., 2022; Lukić et al., 2023). In this study, heat stress during vegetative growth enhanced TPC and TFC in seeds, likely through ROS-induced activation of the phenylpropanoid pathway (Yang et al., 2017; Liu et al., 2021). Besides their role in antioxidant defense, the accumulation of phenolic compounds during seed development is also essential for seed color formation, which is another important component of oil quality (Yu et al., 2014). Sequential cold–heat treatment led to intermediate TPC and TFC levels—higher than cold alone, but lower than heat alone. Antioxidant assays revealed greater free radical scavenging activity (lower DPPH IC_50_) under the combined treatment and the FRAP activity was highest under sequential stress, suggesting additive effects on redox potential. Additionally, the correlations between TPC, TFC and the antioxidant activities were not very strong. Taken together, a diverse range of antioxidants besides phenolics and flavonoids such as ascorbate may contribute to the antioxidant activity under sequential cold-heat stresses (Terpinc et al., 2012; van Buer et al., 2016).

Enzymatic antioxidant responses varied; CAT activity was lowest under cold stress and showed a non-significant decrease under heat and combined treatments. In contrast, SOD activity peaked under sequential cold–heat stress (*p < 0.05*), indicating a shift in ROS detoxification mechanisms. Despite these variations, H_2_O_2_ levels remained stable, supporting the idea of a primed antioxidant network maintaining ROS homeostasis. This is consistent with previous work showing that antioxidant enzyme activity supports cross-tolerance and seedling vigor through priming under temperature stress (Mei et al., 2010). This supports a model of intergenerational stress memory, where early environmental signals shape offspring seed metabolism via integrated regulation of enzymatic and non-enzymatic antioxidant systems, which are sustained through mitosis and meiosis (Lämke & Bäurle, 2017; Friedrich et al., 2019).

Our findings demonstrate that cold stress during the vegetative phase induces significant alterations in seed oil metabolism in rapeseed, characterized by increased total seed oil content and modified fatty acid composition. These changes are accompanied by coordinated transcriptional regulation of key genes involved in fatty acid biosynthesis. The observed upregulation of lipid metabolism under cold-only stress aligns with previous reports (Byun et al., 2014; Çağlı et al., 2025). The cold-induced enhancement of seed oil content likely reflects a metabolic adaptation that reallocates carbon toward lipid biosynthesis, thereby supporting seed energy reserves under stress conditions (Gao et al., 2024).

Notably, cold stress led to a decrease in oleic acid-a monounsaturated fatty acid and an increase in polyunsaturated fatty acids (PUFAs), suggesting a remodeling of seed oil unsaturation to maintain fluidity and oxidative stability—an established strategy in plant stress adaptation (Miquel et al., 1993; Gao et al., 2024). In contrast, heat stress alone or sequential cold–heat treatments did not significantly alter total oil content, indicating that heat may mitigate cold-induced lipid accumulation or activate distinct metabolic pathways. These findings highlight a complex interplay between stress types, where the plant’s acclimation or adaptation strategy likely depends on stress intensity and order of occurrence (Mittler et al., 2022).

Unlike seeds, vegetative tissues typically accumulate low levels of triacylglycerols (TAGs) under normal conditions. However, environmental stress can substantially elevate TAG content in these tissues through transcriptional regulation of fatty acid synthesis genes (Lee et al., 2019; Nam et al., 2022; Zhang et al., 2025). In our study, several fatty acid synthesis genes—including the ω-6 desaturase *BnaFAD2* (involved in converting oleic acid to linoleic acid), *BnaFAD5*, the acyl-ACP thioesterase *BnaFATB*, and *BnaWD40*, a regulatory gene associated with abiotic stress responses—were particularly upregulated under cold stress. These expression patterns suggest that transcriptional responses initiated during the vegetative phase may persist into the reproductive phase, potentially influencing seed development (Dikšaitytė et al., 2019). Further investigation is needed to determine whether such transcriptional memory exists and whether epigenetic mechanisms contribute to this regulatory continuity (Çağlı et al., 2025).

Germination efficiency of progeny was not significantly affected by parental exposure to cold, heat, or sequential cold–heat stress during vegetative growth, with all treatment groups maintaining consistently high germination rates (∼95%). This indicates that the applied stresses did not impair seed viability or embryo development, suggesting that any observed intergenerational effects are unlikely to result from compromised seed quality at the germination stage. Interestingly, seedlings derived from parents exposed to sequential cold and heat stress exhibited significantly higher levels of chlorophyll *a*, chlorophyll *b*, and total chlorophyll compared to all other groups. In contrast, cold stress alone caused a slight, non-significant reduction in chlorophyll content relative to the control. The elevated chlorophyll levels in progeny of sequentially stressed plants imply enhanced photosynthetic capacity, a trait often associated with improved stress resilience and growth vigor (Lukić et al., 2023). The maintenance or enhancement of photosynthetic pigments is a well-documented component of cross-tolerance mechanisms, enabling plants to optimize light capture and sustain carbon assimilation under subsequent abiotic stress.

Consistent with these findings, progeny from sequentially stressed parents also displayed increased seedling fresh weight and higher leaf relative water content (RWC)—physiological traits indicative of greater vigor and improved water retention capacity. These enhancements likely reflect a priming effect inherited from the parental generation, in which stress-induced signaling activates epigenetic or metabolic memory that influences chloroplast development and pigment biosynthesis in offspring (Lämke & Bäurle, 2017; Friedrich et al., 2019). Improved photosynthetic efficiency may also provide the necessary energy and reducing power to support antioxidant defenses and broader metabolic adjustments during stress encounters.

This form of inter- or transgenerational plasticity is supported by numerous studies showing that parental exposure to abiotic stress can prime offspring for enhanced antioxidant defense, osmoprotection, and photosynthetic performance. Accordingly, our results align with previous reports demonstrating that transgenerational stress memory can improve photosynthetic capacity and seedling vigor in progeny (Wang et al., 2016; Hatzig et al., 2018; Lukić et al., 2023). Further investigation is needed to elucidate the specific epigenetic mechanisms underlying this intergenerational stress memory and to clarify how these processes contribute to priming and cross-tolerance in *Brassica napus*.

## Conclusion

While the individual effects of cold acclimation and heat stress in plants are well documented, the intergenerational consequences of sequential cold and heat stress remain underexplored. This study provides evidence that early cold exposure primes *Brassica napus* for enhanced tolerance to subsequent heat stress, with physiological, biochemical, and molecular effects extending into the next, stress-free generation. By examining the progeny of stressed plants, we demonstrate that sequential cold–heat stress induces a distinct and heritable response involving a coordinated regulation at the molecular, biochemical and physiological levels. Notably, progeny from sequentially stressed parents exhibited enhanced photosynthetic capacity, increased vigor, and optimized seed composition—hallmarks of intergenerational stress memory.

These findings highlight the complex cross-talk between temperature stress responses and suggest that priming through early cold exposure can confer cross-tolerance to later heat stress. Elucidating the molecular basis of this intergenerational plasticity will be essential for leveraging stress memory in crop improvement programs aimed at enhancing resilience under increasingly variable climatic conditions.

## Acknowledgments

We would like to thank Murat Yaşar for his help with planting and tending the plants in the greenhouse.

## Author Contribution Statement

Conceptualization: CS; Methodology: IC, AGT, MBH, FT, CS; Software: IC; Investigation: IC, AGT, MBH, FT, CS; Supervision: CS; Funding acquisition: CS; Project administration: CS; Writing - original draft: IC, AGT, CS; Writing - review & editing: IC, CS

## Data Availability Statement

The data that support the findings of this study are available on request from the corresponding author. The data are not publicly available due to privacy or ethical restrictions.

## Conflict of Interest Statement

Authors declare no conflict of interest

## Supplemantary Information

**Table S1.**
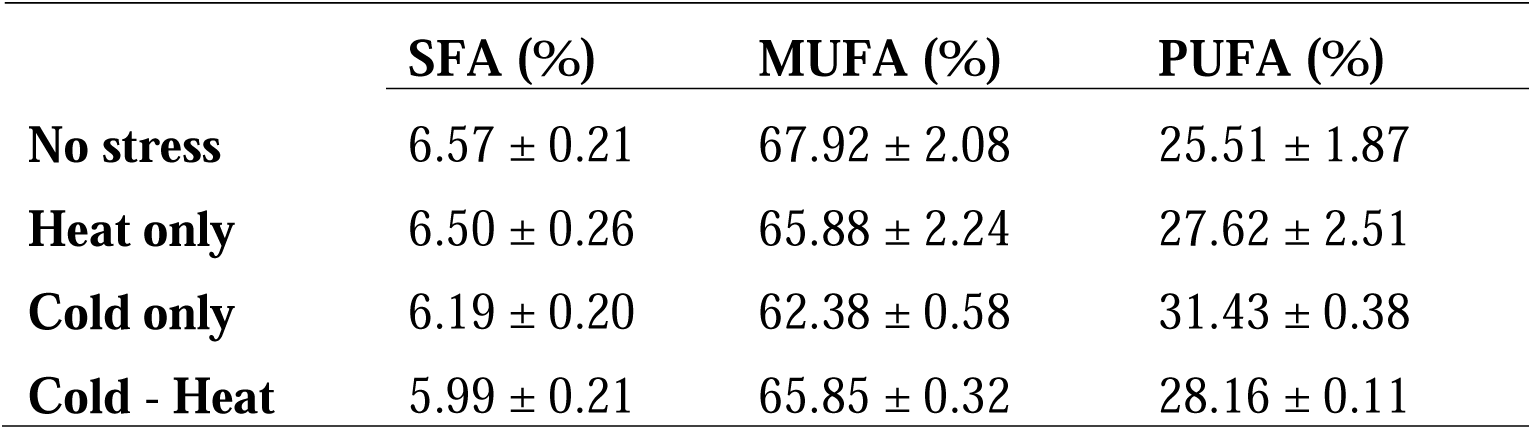
Saturated (SFA), Monounsaturated (MUFA) and Polyunsaturated (PUFA) Fatty Acid Contents of B. napus seeds under different treatments.

**Table S2.** Fresh Seedling Weight of first stress-free generation of rapeseed (*Brassica napus*) under different treatment conditions.

|  | Total fresh seedling weight (g) | Average seedling weight per one seedling (g) |
| --- | --- | --- |
| No stress | 3.180 ± 0.080 | 0.353 ± 0.009 |
| Heat only | 3.370 ± 0.331 | 0.374 ± 0.037 |
| Cold only | 3.300 ± 0.281 | 0.367 ± 0.031 |
| Cold - Heat | 3.765 ± 0.015 | 0.418 ± 0.002 |

Values are means of two biological replicates. Data are represented as Mean ± SEM.

**Table S3.** Leaf Relative Water Content (LRWC) of first stress-free generation of rapeseed (*Brassica napus*) seedlings under different treatment conditions.

|  | LRWC (%) |
| --- | --- |
| No stress | 80.27 ± 0.97 |
| Heat only | 79.82 ± 3.17 |
| Cold only | 85.22 ± 1.47 |
| Cold - Heat | 90.37 ± 0.78 |

Values are represented as Mean ± SEM.

## Notes

### Competing Interest Statement

The authors have declared no competing interest.

## References

Abelenda, J. A., Trabanco, N., Del Olmo, I., Pozas, J., Martín-Trillo, M. D. M., Gómez-Garrido, J., … & Piñeiro, M. (2023). High ambient temperature impacts on flowering time in Brassica napus through both H2A.Z-dependent and independent mechanisms. Plant, Cell & Environment, 46(5), 1427–1441.

Almeselmani, M., Deshmukh, P. S., Sairam, R. K., Kushwaha, S. R., & Singh, T. P. (2006). Protective role of antioxidant enzymes under high temperature stress. Plant Science, 171(3), 382–388.

Arnon, D.I., 1949. Copper enzymes in isolated chloroplasts polyphenol oxidase in Beta vulgaris. Plant Physiology., 24, 1–15.

Baux, A., T. Hebeisen, and D. Pellet. 2008. “Effects of Minimal Temperatures on Low-Linolenic Rapeseed Oil Fatty-Acid Composition.” European Journal of Agronomy 29, no. 2–3: 102–107.

Baux, A., Colbach, N., Allirand, J. M., Jullien, A., Ney, B., & Pellet, D. (2013). Insights into temperature effects on the fatty acid composition of oilseed rape varieties. European Journal of Agronomy, 49, 12–19.

Benitez-Alfonso, Y., Soanes, B. K., Zimba, S., Sinanaj, B., German, L., Sharma, V., … & Foyer, C. H. (2023). Enhancing climate change resilience in agricultural crops. Current Biology, 33(23), R1246–R1261.

Brown, J., Erickson, D. A., Davis, J. B., & Brown, A. P. (1995). Effects of swathing on yield and quality of spring canola (*Brassica napus* L.) in the Pacific Northwest. In Proceedings of the 9th International Rapeseed Congress (Vol. 1, pp. 339–341). Cambridge, UK.

Byun, Y. J., Koo, M. Y., Joo, H. J., Ha-Lee, Y. M., & Lee, D. H. (2014). Comparative analysis of gene expression under cold acclimation, deacclimation and reacclimation in *Arabidopsis*. Physiologia Plantarum, 152(2), 256–274.

Çağlı, İ., Kıvrak, B. E., Altunbaş, O., & Sönmez, Ç. (2025). Unveiling the impact of vernalisation on seed oil content and fatty acid composition in rapeseed (*Brassica napus* L.) through simulated shorter winters. Journal of Agronomy and Crop Science, 211(3), e70057.

Chen, S., Stefanova, K., Siddique, K. H., & Cowling, W. A. (2021). Transient daily heat stress during the early reproductive phase disrupts pod and seed development in *Brassica napus* L. Food and Energy Security, 10(1), e262.

Chen, X., Liu, L., Cai, H., Zheng, B., & Li, J. (2024). Effects of spring low-temperature stress on winter wheat seed-setting characteristics of spike. *Plant*, Soil & Environment, 70(2).

de Almeida, L. M. M., Avice, J. C., Morvan-Bertrand, A., Wagner, M. H., González-Centeno, M. R., Teissedre, P. L., … & Brunel-Muguet, S. (2021). High temperature patterns at the onset of seed maturation determine seed yield and quality in oilseed rape (*Brassica napus* L.) in relation to sulphur nutrition. Environmental and Experimental Botany, 185, 104400.

De Lucia, F., Crevillen, P., Jones, A. M., Greb, T., & Dean, C. (2008). A PHD-polycomb repressive complex 2 triggers the epigenetic silencing of FLC during vernalization. Proceedings of the National Academy of Sciences, 105(44), 16831–16836.

Del Olmo, I., Poza-Viejo, L., Piñeiro, M., Jarillo, J. A., & Crevillén, P. (2019). High ambient temperature leads to reduced FT expression and delayed flowering in *Brassica rapa* via a mechanism associated with H2A.Z dynamics. The Plant Journal, 100(2), 343–356.

Dikšaitytė, A., Viršilė, A., Žaltauskaitė, J., Januškaitienė, I., & Juozapaitienė, G. (2019). Growth and photosynthetic responses in *Brassica napus* differ during stress and recovery periods when exposed to combined heat, drought and elevated CO2. Plant Physiology and Biochemistry, 142, 59–72.

Distéfano, A. M., Bauer, V., Cascallares, M., López, G. A., Fiol, D. F., Zabaleta, E., & Pagnussat, G. C. (2025). Heat stress in plants: sensing, signalling, and ferroptosis. Journal of Experimental Botany, 76(5), 1357–1369.

Elfalleh, W., Nasri, N., Marzougui, N., Thabti, I., M’rabet, A., Yahya, Y., … & Ferchichi, A. (2009). Physico-chemical properties and DPPH-ABTS scavenging activity of some local pomegranate (*Punica granatum*) ecotypes. International Journal of Food Sciences and Nutrition, 60(sup2), 197–210.

Eynck, C., Koopmann, B., Karlovsky, P., & Von Tiedemann, A. (2009). Internal resistance in winter oilseed rape inhibits systemic spread of the vascular pathogen *Verticillium longisporum*. Phytopathology, 99(7), 802–811.

Friedrich, T., Faivre, L., Bäurle, I., & Schubert, D. (2019). Chromatin based mechanisms of temperature memory in plants. *Plant*, Cell & Environment, 42(3), 762–770.

Fu, P., Wilen, R. W., Robertson, A. J., Low, N. H., Tyler, R. T., & Gusta, L. V. (1998). Heat tolerance of cold acclimated puma winter rye seedlings and the effect of a heat shock on freezing tolerance. Plant and Cell Physiology, 39(9), 942–949.

Gao, L., Jiang, H., Li, M., Wang, D., Xiang, H., Zeng, R., … & Shi, Y. (2024). Genetic and lipidomic analyses reveal the key role of lipid metabolism for cold tolerance in maize. Journal of Genetics and Genomics, 51(3), 326–337.

Groot, M. P., Kubisch, A., Ouborg, N. J., Pagel, J., Schmid, K. J., Vergeer, P., & Lampei, C. (2017). Transgenerational effects of mild heat in *Arabidopsis thaliana* show strong genotype specificity that is explained by climate at origin. New Phytologist, 215(3), 1221–1234.

Guo, X., Liu, D., & Chong, K. (2018). Cold signaling in plants: insights into mechanisms and regulation. Journal of Integrative Plant Biology, 60(9), 745–756.

Hatzig, S. V., Nuppenau, J. N., Snowdon, R. J., & Schießl, S. V. (2018). Drought stress has transgenerational effects on seeds and seedlings in winter oilseed rape (*Brassica napus* L.). BMC Plant Biology, 18(1), 297.

Henderson, I. R., Shindo, C., & Dean, C. (2003). The need for winter in the switch to flowering. Annual Review of Genetics, 37(1), 371–392.

Hossain, M. A., Bhattacharjee, S., Armin, S. M., Qian, P., Xin, W., Li, H. Y., … & Tran, L. S. P. (2015). Hydrogen peroxide priming modulates abiotic oxidative stress tolerance: insights from ROS detoxification and scavenging. Frontiers in Plant Science, 6, 420.

Hossain, M. A., Li, Z. G., Hoque, T. S., Burritt, D. J., Fujita, M., & Munné-Bosch, S. (2018). Heat or cold priming-induced cross-tolerance to abiotic stresses in plants: key regulators and possible mechanisms. Protoplasma, 255(1), 399–412.

Huang, R., Liu, Z., Xing, M., Yang, Y., Wu, X., Liu, H., & Liang, W. (2019). Heat stress suppresses *Brassica napus* seed oil accumulation by inhibition of photosynthesis and BnWRI1 pathway. Plant and Cell Physiology, 60(7), 1457–1470.

Jørgensen, R. B., & Andersen, B. (1994). Spontaneous hybridization between oilseed rape (*Brassica napus*) and weedy B. campestris (*Brassicaceae*): a risk of growing genetically modified oilseed rape. American Journal of Botany, 1620–1626.

Kan, Y., Mu, X. R., Gao, J., Lin, H. X., & Lin, Y. (2023). The molecular basis of heat stress responses in plants. Molecular Plant, 16(10), 1612–1634.

Lämke, J., & Bäurle, I. (2017). Epigenetic and chromatin-based mechanisms in environmental stress adaptation and stress memory in plants. Genome Biology, 18(1), 124.

Lee, H. G., Park, M. E., Park, B. Y., Kim, H. U., & Seo, P. J. (2019). The *Arabidopsis* MYB96 transcription factor mediates ABA-dependent triacylglycerol accumulation in vegetative tissues under drought stress conditions. Plants, 8(9), 296.

Li, X., Cai, J., Liu, F., Dai, T., Cao, W., & Jiang, D. (2014). Cold priming drives the sub-cellular antioxidant systems to protect photosynthetic electron transport against subsequent low temperature stress in winter wheat. Plant Physiology and Biochemistry, 82, 34–43.

Liu, B., Zhang, L., Rusalepp, L., Kaurilind, E., Sulaiman, H. Y., Püssa, T., & Niinemets, Ü. (2021). Heat priming improved heat tolerance of photosynthesis, enhanced terpenoid and benzenoid emission and phenolics accumulation in *Achillea millefolium*. Plant, Cell & Environment, 44(7), 2365–2385.

Liu, H., Able, A. J., & Able, J. A. (2022). Priming crops for the future: rewiring stress memory. Trends in Plant Science, 27(7), 699–716.

Lukić, N., Schurr, F. M., Trifković, T., Kukavica, B., & Walter, J. (2023). Transgenerational stress memory in plants is mediated by upregulation of the antioxidative system. Environmental and Experimental Botany, 205, 105129.

Malhotra, H., Sangha, M. K., Pathak, D., Choudhary, O., Kumar, P., & Rathore, P. (2014). A simple hydroponic variant for screening cotton genotypes for salinity tolerance. Crop Improv, 41(2), 134–9.

Mei, Y. Q., & Song, S. Q. (2010). Response to temperature stress of reactive oxygen species scavenging enzymes in the cross-tolerance of barley seed germination. Journal of Zhejiang University SCIENCE B, 11(12), 965–972.

Menon, G., Schulten, A., Dean, C., & Howard, M. (2021). Digital paradigm for Polycomb epigenetic switching and memory. Current Opinion in Plant Biology, 61, 102012.

Miquel, M., James Jr, D., Dooner, H., & Browse, J. (1993). *Arabidopsis* requires polyunsaturated lipids for low-temperature survival. Proceedings of the National Academy of Sciences, 90(13), 6208–6212.

Mittler, R. (2022, November). Developing Climate-Resilient Crops: Improving Plant Tolerance to Multifactorial Stress Combination. *In* ASA, CSSA, SSSA International Annual Meeting. ASA-CSSA-SSSA.

Nam, J. W., Lee, H. G., Do, H., Kim, H. U., & Seo, P. J. (2022). Transcriptional regulation of triacylglycerol accumulation in plants under environmental stress conditions. Journal of experimental botany, 73(9), 2905–2917.

Oberkofler, V., Pratx, L., & Bäurle, I. (2021). Epigenetic regulation of abiotic stress memory: maintaining the good things while they last. Current Opinion in Plant Biology, 61, 102007.

Önder, S., Güvercin, D., & Tonguç, M. (2020). Determination of hydrogen peroxide content and antioxidant enzyme activities in safflower (*Carthamus tinctorius* L.) seeds after accelerated aging test. Süleyman Demirel Üniversitesi Fen Bilimleri Enstitüsü Dergisi, 24(3), 681–688.

Prieto, J. M. (2012). Procedure: Preparation of DPPH Radical, and antioxidant scavenging assay. DPPH Microplate Protocol, 7–9.

Rahaman, M., Mamidi, S., & Rahman, M. (2018). Genome-wide association study of heat stress-tolerance traits in spring-type *Brassica napus* L. under controlled conditions. The Crop Journal, 6(2), 115–125.

Rashid, M., Hampton, J. G., Shaw, M. L., Rolston, M. P., Khan, K. M., & Saville, D. J. (2020). Oxidative damage in forage rape (*Brassica napus* L.) seeds following heat stress during seed development. Journal of Agronomy and Crop Science, 206(1), 101–117.

Secchi, M. A., Fernandez, J. A., Stamm, M. J., Durrett, T., Prasad, P. V., Messina, C. D., & Ciampitti, I. A. (2023). Effects of heat and drought on canola (*Brassica napus* L.) yield, oil, and protein: *A meta-analysis*. Field Crops Research, 293, 108848.

Suter, L., & Widmer, A. (2013). Environmental heat and salt stress induce transgenerational phenotypic changes in *Arabidopsis thaliana*. PloS One, 8(4), e60364.

Terpinc, P., Čeh, B., Ulrih, N. P., & Abramovič, H. (2012). Studies of the correlation between antioxidant properties and the total phenolic content of different oil cake extracts. Industrial crops and products, 39, 210–217.

Thomashow, M. F. (1999). Plant cold acclimation: freezing tolerance genes and regulatory mechanisms. Annual Review of Plant Biology, 50(1), 571–599.

van Buer, J., Cvetkovic, J., & Baier, M. (2016). Cold regulation of plastid ascorbate peroxidases serves as a priming hub controlling ROS signaling in *Arabidopsis thaliana*. BMC Plant Biology, 16(1), 163.

Velikova, V., Yordanov, I., & Edreva, A. (2000). Oxidative stress and some antioxidant systems in acid rain-treated bean plants: protective role of exogenous polyamines. Plant Science, 151(1), 59–66.

Wan, S. B., Tian, L., Tian, R. R., Pan, Q. H., Zhan, J. C., Wen, P. F., … & Huang, W. D. (2009). Involvement of phospholipase D in the low temperature acclimation-induced thermotolerance in grape berry. Plant Physiology and Biochemistry, 47(6), 504–510.

Wang, X., Xin, C., Cai, J., Zhou, Q., Dai, T., Cao, W., & Jiang, D. (2016). Heat priming induces trans-generational tolerance to high temperature stress in wheat. Frontiers in Plant Science, 7, 501.

Whittle, C. A., Otto, S. P., Johnston, M. O., & Krochko, J. E. (2009). Adaptive epigenetic memory of ancestral temperature regime in *Arabidopsis thaliana*. Botany, 87(6), 650–657.

Xiao, F., Xu, T., Lu, B., & Liu, R. (2020). Guidelines for antioxidant assays for food components. Food Frontiers, 1(1), 60–69.

Yang, R., Guo, L., Wang, J., Wang, Z., & Gu, Z. (2017). Heat shock enhances isothiocyanate formation and antioxidant capacity of cabbage sprouts. Journal of Food Processing and Preservation, 41(4), e13034.

Young, L. W., Wilen, R. W., & Bonham-Smith, P. C. (2004). High temperature stress of *Brassica napus* during flowering reduces micro- and megagametophyte fertility, induces fruit abortion, and disrupts seed production. Journal of Experimental Botany, 55(396), 485–495.

Yu, E., Fan, C., Yang, Q., Li, X., Wan, B., Dong, Y., … & Zhou, Y. (2014). Identification of heat responsive genes in *Brassica napus* siliques at the seed-filling stage through transcriptional profiling. PloS One, 9(7), e101914.

Zajac, T., Klimek-Kopyra, A., Oleksy, A., Lorenc-Kozik, A., & Ratajczak, K. (2016). Analysis of yield and plant traits of oilseed rape (*Brassica napus* L.) cultivated in temperate region in light of the possibilities of sowing in arid areas. Acta Agrobotanica, 69(4).

Zhang, J., Venables, I., Callahan, D. L., Zwart, A. B., Passioura, J., Liu, Q., … & Estavillo, G. M. (2025). Water stress enhances triacylglycerol accumulation via different mechanisms in wild-type and transgenic high-leaf oil tobacco. Plant Physiology, 198(1), kiaf151.

